# Spatial heterogeneity in biofilm metabolism elicited by local control of phenazine methylation

**DOI:** 10.1101/2023.02.15.528762

**Authors:** Christopher R. Evans, Marina K. Smiley, Sean Asahara Thio, Mian Wei, Alexa Price-Whelan, Wei Min, Lars E.P. Dietrich

**Author notes:** C.R.E. and M.K.S. contributed equally to this work. Corresponding author: Lars E.P. Dietrich **Email**. **Author contributions**: C.R.E., M.K.S., M.W., and L.E.P.D. designed research; C.R.E., M.K.S., S.A.T., and M.W. performed research; C.R.E., M.K.S., S.A.T., M.W., A.P.-W. and L.E.P.D. analyzed data; W.M. contributed reagents, analytic tools, and expertise; C.R.E., M.K.S, A.P.-W. and L.E.P.D. prepared the manuscript. **Competing Interest Statement**: The authors declare no competing interest. **Classification**: Biological Sciences (major), Microbiology (minor).

## Abstract

Within biofilms, gradients of electron acceptors such as oxygen stimulate the formation of physiological subpopulations. This heterogeneity can enable cross-feeding and promote drug resilience, features of the multicellular lifestyle that make biofilm-based infections difficult to treat. The pathogenic bacterium *Pseudomonas aeruginosa* produces pigments called phenazines that can support metabolic activity in hypoxic/anoxic biofilm subzones, but these compounds also include methylated derivatives that are toxic to their producer under some conditions. Here, we uncover roles for the global regulators RpoS and Hfq/Crc in controlling the beneficial and detrimental effects of methylated phenazines in biofilms. Our results indicate that RpoS controls phenazine methylation by modulating activity of the carbon catabolite repression pathway, in which the Hfq/Crc complex inhibits translation of the phenazine methyltransferase PhzM. We find that RpoS indirectly inhibits expression of CrcZ, a small RNA that binds to and sequesters Hfq/Crc, specifically in the oxic subzone of *P. aeruginosa* biofilms. Deletion of *rpoS* or *crc* therefore leads to overproduction of methylated phenazines, which we show leads to increased metabolic activity—an apparent beneficial effect—in hypoxic/anoxic subpopulations within biofilms. However, we also find that biofilms lacking Crc show increased sensitivity to an exogenously added methylated phenazine, indicating that the increased metabolic activity in this mutant comes at a cost. Together, these results suggest that complex regulation of PhzM allows *P. aeruginosa* to simultaneously exploit the benefits and limit the toxic effects of methylated phenazines.

**Significance Statement:** *P. aeruginosa* causes biofilm-based infections and is known for its production of colorful phenazine derivatives. Among these the methylated phenazines are the most toxic and can cause condition-dependent damage to their producer. In this study, we show that methylated phenazines also have a beneficial effect in that they specifically support metabolic activity at depth in *P. aeruginosa* biofilms, where oxygen limitation would otherwise stall metabolism. We describe a new link between *P. aeruginosa* global regulators that control methylated phenazine production in a manner that limits their toxicity while simultaneously enabling their contribution to metabolism. These results expand our understanding of the strategies that enable *P. aeruginosa* survival in multicellular structures, which is key to its success during chronic host colonization.

## Introduction

Cellular growth at high density promotes the formation of resource gradients, and multicellular structures therefore contain microenvironments with distinct conditions. To maintain metabolic activity, cells within multicellular structures undergo physiological differentiation, which enhances robustness of the overall population (1–5). Co-existence in a multicellular structure also allows differentiated subpopulations to interact in beneficial ways, for example by exchanging metabolites that are optimally produced in one region and consumed in another. The physiological heterogeneity of multicellular structures is problematic in infections or cancers, where the effectiveness of a drug often depends on the metabolic statuses of target cells (6). Approaches that provide insight into physiological differentiation while retaining information about structure can help us to understand the unique biology of multicellularity and can also inform treatment methods.

Microbial biofilms are multicellular structures that exhibit metabolic differentiation and developmental responses to resource limitation, with parallels in metazoans and plants (2, 7). Studies in diverse bacteria have indicated that sigma factors, which are regulatory proteins that mediate transcriptional responses to environmental and physiological cues, are key players in biofilm biology (8–13). The sigma factor is a component of the RNA polymerase that confers specificity for target sequences in promoters, and bacterial genomes typically encode more than one sigma factor with different specificities (8, 14). Thus, condition-dependent shifts in the sigma factor pool allow bacteria to elicit global changes in their transcriptional profiles, which adapt their physiology to their current environment. One of the best-studied sigma factors is RpoS, which is conserved in proteobacteria and typically activates gene expression in stationary phase or during resource limitation (15–18).

The bacterium *Pseudomonas aeruginosa* is a prevalent cause of biofilm-based infections and a model for the study of biofilm development and metabolism. It can adapt to environmental changes over the long term, with its high mutation rate (19–22), and in the short term, with an arsenal of up to 26 sigma factors (depending on strain identity) and hundreds of transcription factors (23–25). These regulators form a complex network of gene expression control that we are only beginning to understand holistically (24, 25). Our knowledge about the roles of sigma factor-controlled networks in regulating gene expression and affecting *P. aeruginosa* physiology is particularly limited for cells surviving in more complex environments like biofilms.

RpoS is required for wild-type *P. aeruginosa* biofilm development in a variety of assays and across strain backgrounds. It is expressed in biofilms, particularly at the biofilm-air interface, in *Escherichia coli* and *P. aeruginosa* and its regulon includes genes involved in crucial redox metabolism in the oxic biofilm subzone (10, 26–28). Furthermore, *rpoS* mutants have been reported to overproduce pyocyanin (1-hydroxy-5-methylphenazine) (29), one of several phenazine compounds that allow *P. aeruginosa* to carry out extracellular electron transfer when respiratory electron acceptors are limiting (such as within biofilms) (30–32). While studies in biofilms have indicated that this electron-shuttling property of phenazines supports metabolic activity, recent work has also shown that phenazines can be toxic to *P. aeruginosa* under specific conditions or when their production and localization are not properly controlled (33–35).

In this study, we examined the role of RpoS in *P. aeruginosa* biofilm physiology and were intrigued to find that this sigma factor has an inhibitory effect on metabolic activity at depth in biofilms. At first glance, an inhibition of metabolic activity seems detrimental, and we sought to understand how and why a major regulator would mediate this effect. Here, we describe identification of a regulatory pathway and of phenazine products that act downstream of RpoS to modulate metabolic activity. Our findings implicate this pathway in methylated phenazine resistance and suggest that modulation of phenazine methylation is critical for balancing the toxic and beneficial effects of these reactive compounds in *P. aeruginosa* biofilms.

## Results

### RpoS attenuates biofilm metabolic activity

Prior work has indicated that RpoS is active in biofilms and that it confers fitness during conditions of either carbon source or oxygen limitation in *P. aeruginosa (36, 37)*. To test whether RpoS contributes to metabolic activity in biofilms, we prepared thin sections of 3 day-old *P. aeruginosa* PA14 WT and Δ*rpoS* colony biofilms that had been transferred to medium containing deuterated water for 5 hours (**FIGURE 1A**). The incorporation of deuterium into biomass, as a proxy for metabolic activity, was then imaged by stimulated Raman scattering (SRS) microscopy (31). We found that Δ*rpoS* showed enhanced metabolic activity across biofilm depth compared to WT (**FIGURE 1B**), notably at depths of 70 μm or more. Microelectrode studies have shown that oxygen is depleted below ~75 μm leaving this lower portion of the biofilm hypoxic/anoxic (38).

**Figure 1.**
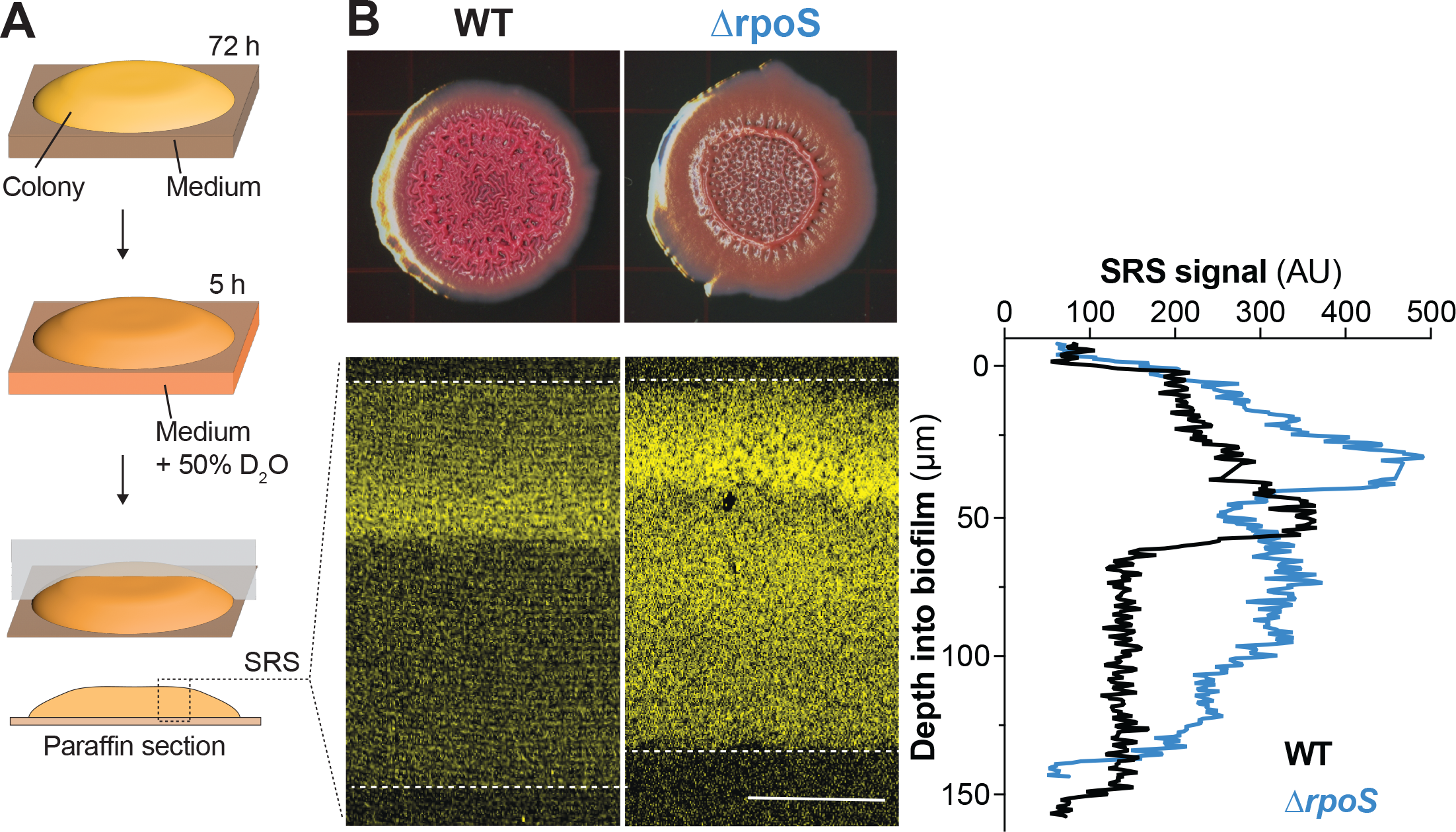
Δ*rpoS* biofilms show increased metabolic activity at depth when compared to those formed by WT PA14. (**A**) Scheme depicting the preparation of colony biofilm thin sections for imaging by stimulated Raman scattering (SRS) microscopy. (**B**) Top: PA14 and Δ*rpoS* biofilms grown for 72 hours on 1% tryptone, 1% agar medium containing the dyes Congo red and Coomassie blue. Bottom: SRS microscopy images of PA14 and Δ*rpoS* biofilm thin sections. The top and bottom of each biofilm is indicated by a dotted line, and the SRS signal is false-colored yellow. Right: quantification of average SRS signal across depth. Experiments were performed in biological triplicate and representative images are shown. Scale bar is 50 μm.

### Methylated phenazines support metabolic activity across depth in PA14 biofilms

We have previously reported that phenazine production (**FIGURE 2A**) enhances metabolic activity at depth in PA14 biofilms (31) and we therefore hypothesized that the effects of RpoS on biofilm metabolic activity could be mediated by effects on phenazine production. To test this, we imaged biofilm thin sections from a Δ*rpoS*Δ*phz* mutant, which is unable to produce phenazines. Removing the capacity to produce phenazines abolished the enhanced metabolic activity of the Δ*rpoS* parent strain, indicating that this phenotype is indeed phenazine-dependent (**FIGURE 2B**).

**Figure 2.**
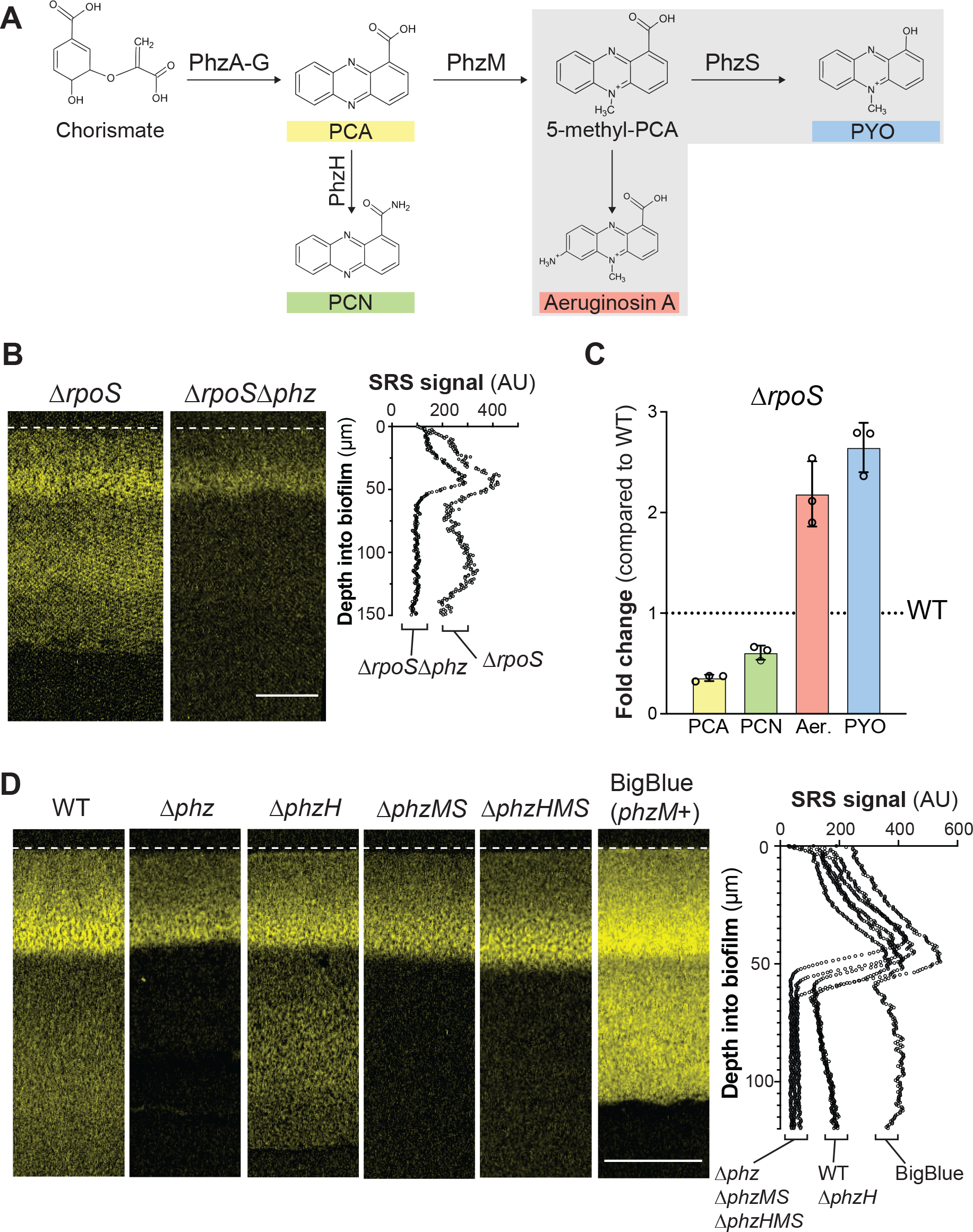
Biofilm metabolic activity at depth is supported specifically by the production of methylated phenazine derivatives. (**A**) The phenazine biosynthetic pathway. A series of biosynthetic enzymes convert two molecules of chorismate to phenazine-1-carboxylic acid (PCA). PCA can be aminated by PhzH to produce phenazine-1-carboxamide (PCN) or methylated by PhzM to produce 5-methyl-PCA. Further modification of 5-methyl-PCA abiotically by free amine groups or enzymatically by PhzS produces aeruginosins (one example structure is shown) or pyocyanin, respectively. (**B**) SRS microscopy images of thin sections prepared from Δ*rpoS* and Δ*rpoS*Δ*phz* biofilms. (**C**) Change in phenazine production by Δ*rpoS* biofilms relative to biofilms formed by WT PA14. Biofilms were grown on 1% tryptone, 1% agar for 72 hours. Aer., aeruginosins. Individual points represent data for biological triplicates and error bars represent standard deviation. (**D**) SRS microscopy images of thin sections prepared from PA14 WT and phenazine biosynthetic mutant biofilms. For (C) and (D), SRS signal is false-colored yellow; quantification of the average SRS signal across depth is shown in the right-hand panels; and experiments were performed in biological triplicate and representative images are shown. Scale bars are 50 μm.

Due to the importance of phenazines in the high metabolic activity of Δ*rpoS* biofilms, we investigated how RpoS affects the production of specific phenazine derivatives. *P. aeruginosa* phenazines are produced in biofilms and in stationary phase during growth in shaken liquid cultures (39). The first derivative produced in the *P. aeruginosa* phenazine synthesis pathway is phenazine-1-carboxylic acid (PCA), which can be converted to (i) phenazine-1-carboxamide (PCN) via the PhzH transamidase, or (ii) 5-methyl-PCA via the PhzM methyltransferase. 5-methyl-PCA, an unstable intermediate, is converted to other methylated phenazines including pyocyanin and aeruginosin A (40–42) (**FIGURE 2A**). It has been reported that RpoS inhibits production of PhzM, and that RpoS-deficient mutants show increased pyocyanin and decreased PCA production (29, 43). HPLC and spectrophotometer analysis of extracts from biofilms and the underlying growth medium confirmed that flux through the PhzM step of the phenazine biosynthetic pathway was increased in Δ*rpoS* biofilms (**FIGURE 2C**). Furthermore, PCA levels in PA14 and Δ*rpoS* biofilms lacking the phenazine modification enzymes PhzH, PhzM, and PhzS (Δ*phzHMS*) were similar, providing further evidence that the changes in phenazine differentiation observed in Δ*rpoS* biofilms is due to changes in regulation of one or more enzyme(s) that modify PCA (**FIGURE S1**).

To evaluate whether specific phenazine derivatives contribute to metabolic activity in biofilms and whether this underpins the Δ*rpoS* phenotype, we incubated mutant biofilms lacking genes for phenazine-modification enzymes on deuterated water, prepared thin sections, and imaged via SRS microscopy. We observed a clear correlation between the production of methylated phenazines and the amount of metabolic activity across depth in biofilms, most notably in the hypoxic/anoxic zone. Consistent with a specific role for methylated phenazines in promoting metabolic activity, a strain (“BigBlue”) that contains a second copy of *phzM* on the chromosome (44) and therefore overproduces methylated phenazines, showed the strongest SRS signal (**FIGURE 2D**).

### RpoS regulates phenazine derivatization via the *P. aeruginosa* carbon catabolite repression system

We sought to further understand the regulatory pathway that allows RpoS to control phenazine methylation. The *phzM* transcript is one of >200 mRNA targets of the Hfq/Crc (“catabolite repression control”) protein complex (**FIGURE 3A**) (45–48). The Hfq/Crc complex mediates the terminal regulatory step of the carbon catabolite repression (CCR) pathway. Canonically, the CCR pathway acts to inhibit translation of mRNAs that support the use of nonpreferred carbon sources when preferred substrates are available (49, 50), though recent studies have indicated that it has broad effects on *P. aeruginosa* physiology beyond primary metabolic functions (51–53). Consistent with the notion that RpoS acts via this pathway to control phenazine methylation, a recent study (43) found that the translation of *phzM* is inhibited by RpoS. To confirm this, we created a translational reporter construct that contains the *phzM* promoter plus the first six codons of *phzM* (therefore including the native ribosomal binding site and the Hfq/Crc binding site (54)), fused to *mScarlet*. Examining expression of this reporter in stationary phase during growth in liquid cultures, we saw similar increases in mScarlet fluorescence when this construct was expressed in the Δ*rpoS* or Δ*crc* backgrounds compared to expression in the WT (**FIGURE 3B**). In further support of this model, we also found that, like Δ*rpoS* biofilms, Δ*crc* and Δ*rpoS*Δ*crc* biofilms show enhanced production of methylated phenazines (**FIGURE 3C**).

**Figure 3.**
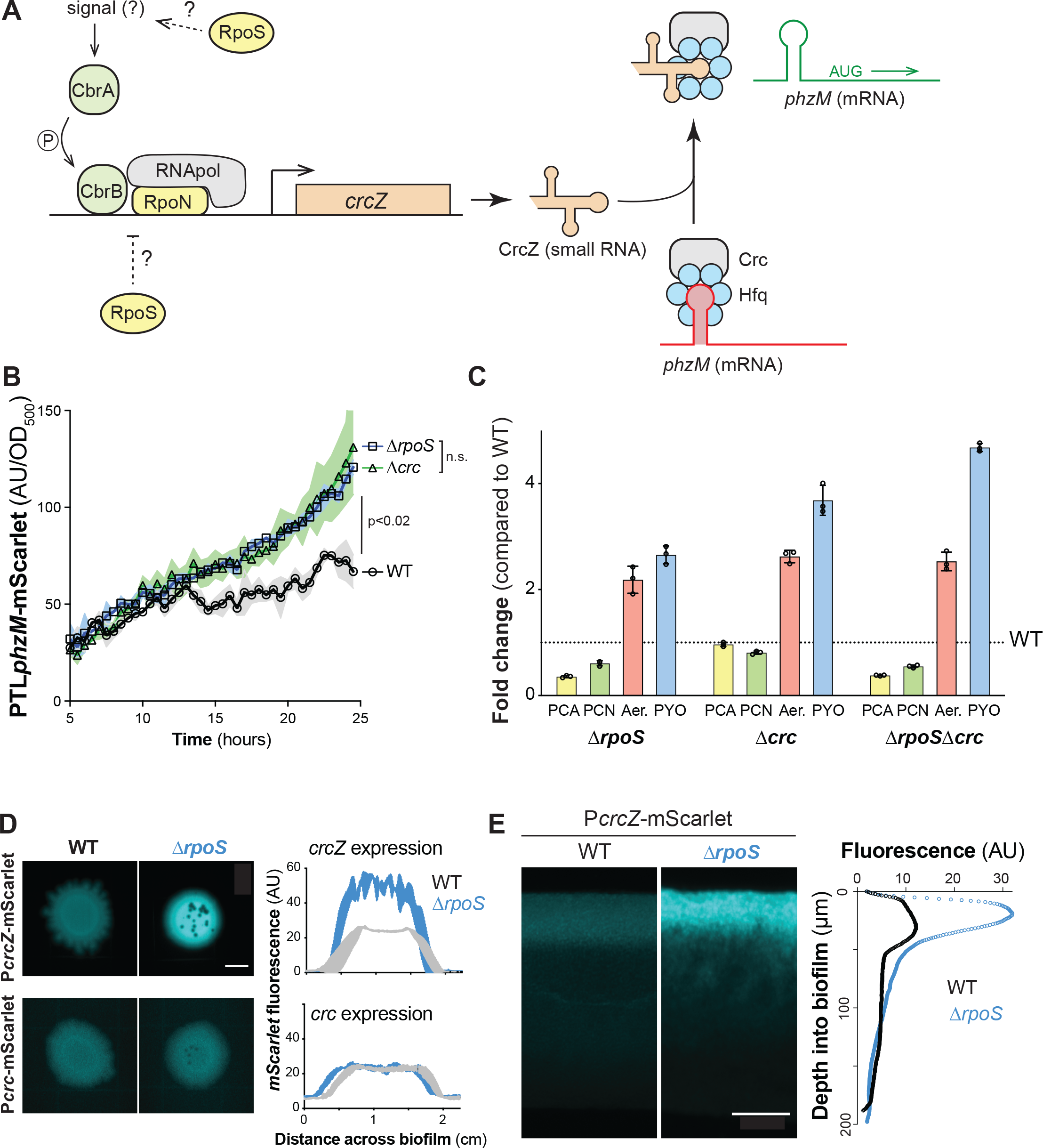
RpoS inhibits phenazine methylation by modulating the carbon catabolite repression pathway at the biofilm-air interface. (**A**) Schematic of the *P. aeruginosa* carbon catabolite repression pathway, which controls translation of *phzM* in a carbon source-dependent manner. RpoN is the canonical sigma factor that is associated with this pathway and it acts in conjunction with the transcription factor CbrB. When CrcZ levels are low, the Crc/Hfq complex binds to *phzM* transcript and inhibits translation. When CrcZ levels are high, CrcZ binds and sequesters Crc/Hfq, allowing translation of the *phzM* transcript. (**B**) Fluorescence conferred by PTL*phzM*-*mScarlet* over time for planktonic cultures of the indicated parent strains. Readings are shown starting at 5 h of incubation, which corresponds to the onset of stationary phase. (**C**) Changes in production of phenazine derivatives by Δ*rpoS*, Δ*crc*, and Δ*rpoS*Δ*crc* biofilms relative to biofilms formed by WT PA14. Individual points represent biological triplicates and error bars represent standard deviation. (**D**) Representative whole-biofilm fluorescence images of PA14 WT and Δ*rpoS* strains expressing mScarlet under control of *PcrcZ* or P*crc*. mScarlet fluorescence is false-colored cyan. The average levels of fluorescence produced by each strain along the diameter of three independent biofilms with the P*crc*-*mScarlet* or P*crcZ*-*mScarlet* reporters are shown in the right-hand plots. Scale bar is 5 mm. (**E**) Fluorescence microscopy of 13 thin sections prepared from WT PA14 and Δ*rpoS* biofilms expressing P*crcZ-mScarlet*. mScarlet fluorescence is false-colored cyan. Average fluorescence across depth is shown in the righthand plot. In (B) and (D), the average of 3 biological replicates is plotted and shading indicates the standard deviation. *p* values were calculated using unpaired, two-tailed *t*-tests. Scale bar is 50 μm.

### RpoS inhibits *crcZ* expression

Binding of the Hfq/Crc complex to target transcripts can be prevented by the small RNA CrcZ (**FIGURE 3A**), which itself can bind and thereby sequester Hfq/Crc under appropriate conditions. This regulatory mechanism allows for condition-dependent translation of Hfq/Crc targets (50). RpoS could therefore affect Hfq/Crc activity by modulating transcription of either *crcZ* or *crc*. To test this, we created constructs in which the upstream promoter regions of *crcZ* or *crc* are cloned in front of *mScarlet* (P*crcZ* and P*crc*, respectively) and then placed these in a neutral site on the chromosome in WT and Δ*rpoS* backgrounds. We found that in both biofilms and liquid (planktonic) cultures, removal of RpoS did not affect activity of P*crc*, but, in contrast, led to a strong enhancement of P*crcZ* activity (**FIGURE 3D, FIGURE S2A**). This indicates that RpoS inhibits *crcZ* expression. Because RpoS is a sigma factor, its binding to promoter sequences acts solely to induce expression and therefore its effect on *crcZ* expression does not arise from direct interaction with the *crcZ* promoter.

The resource gradients that form across depth in colony biofilms lead to physiological differentiation and subpopulations with distinct patterns of gene expression (2, 4, 5, 26, 55). RpoS-dependent effects on gene expression could be localized to a specific biofilm subzone, perhaps where this activity would be most physiologically relevant. Indeed, results from previous studies suggest that RpoS is most expressed and active at the *P. aeruginosa* biofilm-air interface (26, 27, 56, 57). Moreover, pyocyanin synthesis requires oxygen, and prior work by our group has indicated that production of 5-Me-PCA may also be limited to oxygen-containing biofilm subzones (58). When we prepared thin sections of WT and Δ*rpoS* biofilms expressing P*crcZ*-*mScarlet* and imaged by fluorescence microscopy, we found that the indirect effect of RpoS on *crcZ* expression was indeed localized to the biofilm-air interface (**FIGURE 3E**), similar to what has been observed for direct RpoS targets such as the promoter for *coxB* (26, 38). The observation that RpoS inhibits *crcZ* expression specifically at the biofilm-air interface, which is the biofilm region furthest from the growth medium, is consistent with a role for RpoS and the CCR system in inhibiting methylated phenazine production. Because phenazines are excreted small molecules with the potential to diffuse through the biofilm, changes in phenazine derivatization due to changes in RpoS activity at the biofilm surface can negatively impact metabolic activity at depth in biofilms (**FIGURE 1B**).

Evidence suggests *crcZ* transcription in *P. aeruginosa* is driven directly by the regulator CbrB and the sigma factor RpoN **(FIGURE 3A, FIGURE S2A**) (24, 50, 59). CbrB is part of the CbrAB two-component system, which responds to an as-yet-unidentified cue. Our observation that RpoS inhibits *crcZ* transcription raised the question of whether RpoS might be acting upstream of CbrAB. To test this, we deleted *cbrB* in the Δ*rpoS* P*crcZ-mScarlet* reporter strain and found that this abolished P*crcZ* activity (**FIGURE S2B**). These results indicate that CbrB is required for *crcZ* repression in the Δ*rpoS* background, and are consistent with a model in which RpoS acts upstream of CbrAB, the first apparent step of the *P. aeruginosa* CCR pathway, to inhibit *phzM* translation and therefore methylated phenazine production. In other words, RpoS might inhibit CbrB-induced *crcZ* expression, leading to a larger pool of un-sequestered Crc available to bind *phzM* transcript, and therefore lower levels of *phzM* translation in the WT when compared to Δ*rpoS*.

Our observations linking RpoS to the CCR pathway and the inhibition of metabolic activity in biofilms suggest that RpoN, which is required for *crcZ* expression, could promote metabolic activity in biofilms. We addressed this by imaging thin sections of WT and Δ*rpoN* biofilms via SRS microscopy, and found that Δ*rpoN* biofilms showed lower levels of metabolic activity across depth than those formed by the WT (**FIGURE S3**). This observation is consistent with RpoN’s role in the CCR pathway and the abrogation of methylated phenazine production in this mutant (54, 60). A second prediction that arises from the fact that RpoS inhibits *crcZ* expression (**FIGURE 3D**) is that RpoS might hinder *P. aeruginosa*’s ability to transition to growth on non-preferred carbon sources, as *crcZ* relieves Crc/Hfq repression of target mRNAs involved in the use of these substrates (61). To test this idea, we used a defined liquid growth medium with a highly preferred carbon source (i.e., succinate, on which *P. aeruginosa* shows low levels of *crcZ* expression) or with a non-preferred carbon source (i.e., mannitol, which induces high levels of *crcZ* expression) (62). We found that the Δ*rpoS* mutant indeed showed a large increase in *crcZ* expression (**FIGURE S4**), and enhanced growth during the lag and exponential phases, during growth on mannitol when compared to WT (**FIGURE S5**). During growth on succinate, Δ*rpoS* did not show a large increase in *crcZ* expression or enhanced growth during the lag or exponential phases of growth. The Δ*rpoS* mutant did, however, show an increased growth yield relative to WT during growth on succinate, which may arise from independent effects of this mutation.

### Δ*crc* shows increased metabolic activity but is sensitive to phenazine methosulfate

Due to its role in stabilizing Hfq’s interaction with target transcripts such as that of *phzM*, mutants lacking Crc overproduce PhzM and methylated phenazines (54, 63) (**FIGURE 3C**). Having observed that Δ*rpoS* and BigBlue—two mutants with elevated methylated phenazine production—show increased metabolic activity at depth in biofilms we wondered whether this correlation would extend to the Δ*crc* mutant. We used SRS microscopy to image Δ*crc* biofilm thin sections and found that this mutant also showed increased metabolic activity across depth (**FIGURE 4A**). Together, the phenotypes we observed for the Δ*rpoS* (**FIGURE 1B**) and Δ*crc* mutants indicate that RpoS and Hfq/Crc inhibit physiological processes–leading to decreased metabolic activity and growth–raising the question of whether RpoS and Hfq/Crc activity have any benefits for PA14 under the conditions used here. In other words, do RpoS and/or Hfq/Crc have positive impacts that outweigh their inhibition of metabolic activity and growth?

**Figure 4.**
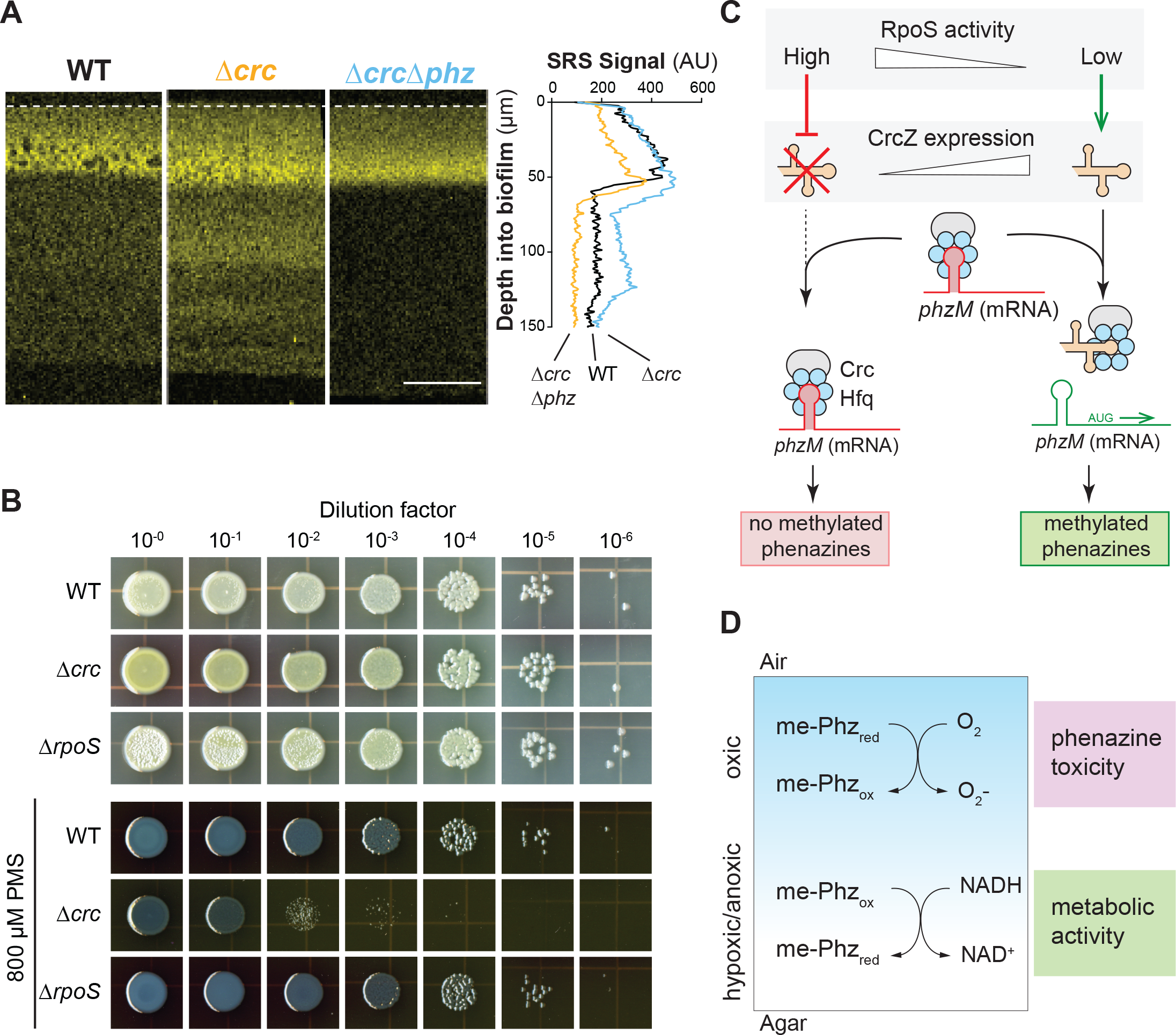
*crc* inhibits phenazine-supported metabolic activity but contributes to methylated phenazine resistance. (**A**) SRS images of thin sections prepared from WT, Δ*crc*, and Δ*crc*Δ*phz* biofilms. SRS signal is false-colored yellow and the average signal across depth is shown in the right-hand plot. Scale bar is 50 μm. (**B**) Growth of WT, Δ*rpoS*, and Δ*crc* on 1% tryptone, 1% agar medium without or with 800 μM PMS. (**C**) Scheme depicting the regulatory relationship between RpoS activity and *phzM* translation. High RpoS activity indirectly inhibits production of the sRNA CrcZ, liberating the Crc/Hfq complex, which in turn represses *phzM* translation. When RpoS activity is low, *crcZ* is transcribed and the CrcZ sRNA binds to and sequesters the Crc/Hfq complex; this allows *phzM* translation to proceed. (**D**) Model depicting the differential effects of methylated phenazines in the oxic and hypoxic/anoxic subzones of a biofilm. We propose that methylated phenazines are toxic in the oxic subzone, due to low nutrient availability and the reactivity of phenazines with oxygen, and that cells in this region benefit from RpoS- and Hfq/Crc-mediated dampening of phenazine methylation. Cells in the hypoxic/anoxic subzone, in contrast, benefit from phenazine methylation because methylated phenazines serve as alternate acceptors of cellular reducing equivalents and support metabolic activity. NADH is shown as a representative carrier of cellular reducing equivalents. The blue shading represents the O_2_ gradient. For (A) and (C), experiments were performed with biological triplicates and representative data are shown.

Previous work has shown that pyocyanin toxicity in *P. aeruginosa* arises specifically under conditions of electron donor/carbon source limitation, which are also conditions that correlate with RpoS activity and which arise at the biofilm-air interface in our colony biofilm model (**FIGURE 3B**) (18, 26, 27, 33). We therefore reasoned that, although methylated phenazine production boosts metabolic activity at depth in biofilms, the net effect of increased methylated phenazine production on the population might be detrimental due to the toxicity of these compounds at the biofilm surface. To test whether RpoS and/or Hfq/Crc mitigates the toxicity of methylated phenazines, we plated the WT and the Δ*crc* and Δ*rpoS* mutants on medium containing the synthetic compound phenazine methosulfate (PMS). PMS is structurally similar to the methylated phenazines produced by *P. aeruginosa* and we have shown that other mechanisms of *P. aeruginosa* self-protection are active against this compound (34, 35). We found that Δ*crc*, but not Δ*rpoS*, showed a greater inhibition of growth by PMS than the WT (**FIGURE 4B**). The Δ*crc* phenotype is in line with previous work in *P. aeruginosa* strain PAO1, which indicated that the Δ*crc* mutation affects redox homeostasis, tolerance of oxidizing agents, and expression of genes involved in the oxidative stress response (64). In the absence of Crc, target mRNAs whose translation would be repressed by Hfq/Crc in the wild-type background are translated. Evidence suggests that these include proteins that carry out steps in the Entner-Doudoroff and pentose phosphate pathways and that their increased levels in Δ*crc* lead to effects on redox balancing. In the Δ*rpoS* background, RpoS-dependent effects on *crcZ* levels are absent but Crc is still present, and it is possible that other regulators modulate *crcZ* levels to allow Crc to contribute to oxidative stress resistance in the presence of PMS.

## Discussion

Besides *P. aeruginosa (65)*, several other pseudomonad species including *P. chlororaphis* and *P. synxantha* make the phenazine PCA (38, 66, 67). However, among these species *P. aeruginosa* is unusual in its ability to produce methylated phenazines (35). Out of the variety of *P. aeruginosa* phenazines, those that are methylated have the most dramatic effects on biofilm matrix production and morphogenesis (34), antibiotic tolerance (31, 68), and—as shown in this study—metabolic activity. In this context it is interesting that, as indicated by our findings, phenazine methylation is linked to global regulatory pathways that respond to carbon source identity and abundance. Phenazine production itself is a carbon- and nitrogen-consuming process (69), but in addition methylated phenazines–which have relatively high redox potentials and reactivities–in particular have the effect of oxidizing the cytoplasm and activating cellular defense mechanisms including the expression of efflux pumps (68, 70–72). Because nutritional conditions affect the cellular carbon/nitrogen balance and could serve as proxy signals for environments in which these mechanisms contribute to fitness, it could be beneficial for *P. aeruginosa* to control phenazine methylation so that it occurs when specific types of carbon sources are available (73).

**FIGURES 4C** and **4D** depict models of the integrated effects of RpoS and the CCR pathway on metabolic activity and survival in PA14 biofilms. The two-component system CbrAB, which functions as the first step in the pseudomonad CCR pathway, is sensitive to carbon source identity but its precise activating signal has not been described. Several studies have suggested that the CbrA-activating signal is intracellular and representative of the carbon/nitrogen balance (50, 62, 74, 75). Once activated, CbrA transfers a phosphoryl group to CbrB, which binds directly to the *crcZ* promoter and recruits RpoN-containing RNA polymerase to stimulate transcription (50). The sRNA CrcZ binds and sequesters Crc/Hfq, allowing translation of *phzM* mRNA and thereby promoting phenazine methylation (45). In WT PA14 biofilms (**FIGURE 1**) metabolic activity is supported by oxygen available at the biofilm-air interface and by methylated phenazines available at depth in the hypoxic/anoxic region. In Δ*rpoS* and Δ*crc* biofilms (**FIGURE 1**) metabolic activity is enhanced throughout the biofilm due to increased production of methylated phenazines. In the Δ*crc* background, this enhanced metabolic activity apparently comes at a cost as evidenced by increased sensitivity to methylated phenazine (**FIGURE 4B**). Experiments in liquid cultures have indicated that the methylated phenazine pyocyanin is toxic to producing cells that are starved for carbon (33). Additionally, Crc has been shown to prevent the accumulation of reactive oxygen species and the activation of oxidative stress pathways (64). This suggests a trade-off may exist in biofilms between the self-preservation of oxic cells and the potential metabolic activity of cells in hypoxic/anoxic regions of the biofilm, balanced by the dampening of phenazine methylation by RpoS acting via the CCR system.

We conclude that RpoS promotes expression of genes that enhance survival during carbon source limitation, but also inhibits expression of *crcZ*, at the biofilm-air interface. The mechanism of this inhibition is unclear but could arise from the effects of the RpoS regulon on *P. aeruginosa* metabolism; this regulon includes genes involved in primary metabolic pathways and RpoS activity could stimulate a shift in the *P. aeruginosa* metabolome that is sensed by CbrAB (18, 24). An alternative explanation could be that RpoS inhibits *crcZ* expression by competing with RpoN for binding to the RNA polymerase. However, previous analysis of sigma factor deletion strains does not show correlative positive associations between expression of RpoS-dependent genes in *rpoN*-deletion strains, nor vice versa (24), suggesting that this is not the mechanism of RpoS-dependent *crcZ* repression. Regardless, we have observed that phenazine methylation is limited by an RpoS-dependent inhibition of *crcZ* expression at the surface of *P. aeruginosa* biofilms. Because methylated phenazines contribute to metabolic activity in the hypoxic/anoxic biofilm subzone, RpoS’ effects “trickle down” to this region. RpoS- and Crc-dependent inhibition of methylated phenazine production in biofilms may therefore represent a physiological tradeoff that limits toxicity at the air-interface at the expense of metabolic activity at depth.

## Materials and Methods

### Strains and Growth Conditions

All bacterial strains used in this study are listed in **TABLE S1**. For routine liquid culture growth, single colonies of *Pseudomonas aeruginosa* UCBPP-PA14 were used to inoculate lysogeny broth (LB) or 1% tryptone, as specified in the text, and shaken at 37°C, 200 rpm. For biofilm growth, strains were grown in liquid LB for 12-16 hours, then diluted 1:100, or 1:50 for Δ*rpoN* backgrounds, in 2 mL LB and grown for 2.5 hours at 37°C to an OD(500 nm) of 0.4-0.6. These liquid cultures were spotted onto 1% tryptone 1% agar plates and incubated at 25°C. Biofilm morphology was determined as described previously (76) on morphology assay plates made of 1% tryptone and 1% agar and containing 20 μg/mL Congo red and 30 μg/mL Coomassie blue (77). Images of colonies were taken using a Keyence VHX-1000 digital microscope.

To compare the growth dynamics and activities of reporters for individual strains, experiments were carried out in 96-well plate, black-sided, clear-bottomed plates (Greiner Bio-One). Strains were pre-grown in glass culture tubes in liquid LB for 12-16 hours, then used inoculate 200 μL of media at a dilution of 1:100 in 96-well plates. Plates were incubated at 37°C (PTL_*phzM*_-*mScarlet*) or 25°C (P_*crcZ*_-*mScarlet*) with continuous shaking on the medium setting in a Biotek Synergy H1 plate reader. The defined medium used to test individual carbon sources was MOPS minimal medium (50 mM 4 morpholinepropanesulfonic acid (pH 7.2), 43 mM NaCl, 93 mM NH_4_Cl, 2.2 mM KH_2_PO_4_, 1 μg/mL FeSO_4_•7H_2_O, 1 mM MgSO_4_•7H_2_O). Growth was assessed by taking OD readings at 500 nm and fluorescence readings at excitation and emission wavelengths of 569 nm and 599 nm respectively every 30 minutes for up to 40 hours.

### Fluorescent Reporter Strain Construction

All plasmids used in this study are listed in **TABLE S2**. To create reporters of promoter activity for the *crcZ* and *crc* genes, the respective 161bp and 500bp promoter regions upstream of the start codon were amplified by PCR using primers (**TABLE S3**) that added KpnI and XhoI (P*crc*) or SpeI and XhoI (P*crcZ*) digest sites to the 5’ and 3’ ends of the sequence. Purified PCR products were digested and ligated into the pSEK109 vector at the multiple cloning site, upstream of the *mScarlet* sequence. The resulting plasmids were transformed into *E. coli* strain UQ950 and verified by sequencing. They were then moved into *P. aeruginosa* wild-type or mutant strains, as indicated in **TABLE S1**, using biparental conjugation with *E. coli* strain S17-1. Selection for recombinants was carried out as previously described (38). To create the reporter of PhzM translation, the upstream region and first six codons of *phzM* was fused to *mScarlet* using nested PCR with primers that added EcoRI and SphI to the 5’ and 3’ ends of the fusion product. The fusion product was digested and ligated into the pSEK103 vector, sequenced, transformed into UQ950, and moved into *P. aeruginosa* strains as described above.

### Creation of Deletion Mutant Strains

All plasmids used in this study are listed in **TABLE S2**. Deletion constructs were made and deletion strains produced via homologous recombination as previously described. Briefly, ~1 kb genomic flanks on either side of the gene to be deleted were amplified via PCR (primers listed in **TABLE S3**) and ligated into digested pMQ30 plasmid using Saccharomyces cerevisiae InvSc1 gap-closing repair. Successful ligations were selected with SD-Ura plates, grown, and the plasmid purified by miniprep. The plasmid was transformed into an *E. coli* mating strain, BW29427, and mated into the recipient *P. aeruginosa* strain. Plasmid integration into *P. aeruginosa* resulted in merodiploids, which are resolved by outgrowth in LB. Strains that successfully lost the integrated plasmid via homologous recombination were selected by growth on sucrose. The final clones were confirmed by PCR with primers flanking the excision site.

### Biofilm Thin Sectioning

Colony preparation and sectioning was performed as previously described with minor modifications (78). Bacterial subcultures were spotted onto 45mL/15mL two-layer 1% tryptone 1% agar in 100 mm x 15 mm square plates and grown for 72 hours. The biofilms were overlaid with 25 mL of 1% molten agar, solidified, excised into histocassettes, and fixed in 4% paraformaldehyde in PBS for 24 hours. After fixing, the samples were washed in a series of increasing ethanol concentrations in PBS, up to 100%, then histoclear, then infiltrated with molten paraffin wax. The samples were then placed in paraffin wax in wax molds and solidified overnight. The wax blocks containing the biofilm samples were shaped and the middle of the biofilms sectioned into 10 μm sections. The sections were floated on water and placed onto pre-charged glass slides. The slides were dried for 48 hours, then heat fixed on a 45°C hot plate for 45 minutes. Once cooled, the samples were rehydrated in histoclear, then a series of decreasing ethanol concentrations in PBS, down to 0% ethanol. Once rehydrated, mounting media and a coverslip was placed on the slide and the samples were imaged.

### Stimulated Raman Scattering Microscopy

Five microliters of a bacterial subculture were spotted onto 45mL/15mL two-layer 1% tryptone 1% agar plates were grown for 72 hours, then the top layer and biofilm was transferred to 2 mL 1% tryptone 1% agar 50% D_2_O and incubated at 25°C for 5 hours. The biofilms were then thin-sectioned as described above. For SRS microscopy, an integrated laser source (picoEMERALD, Applied Physics & Electronics, Inc.) was used to produce both a Stokes beam (1064□nm, 6□ps, intensity modulated at 8□MHz) and a tunable pump beam (720–990□nm, 5– 6□ps) at an 80□MHz repetition rate. The spectral resolution of SRS is FWHM□=□6–7□cm^−1^. These two spatially and temporally overlapped beams with optimized near-IR throughput were coupled into an inverted multiphoton laser-scanning microscope (FV1200MPE, Olympus). Both beams were focused on the biofilm samples through a 25X water objective (XLPlan N, 1.05 N.A. MP, Olympus) and collected with a high N.A. oil condenser lens (1.4 N.A., Olympus) after the sample. By removing the Stokes beam with a high O.D. bandpass filter (890/220 CARS, Chroma Technology), the pump beam is detected with a large area Si photodiode (FDS1010, Thorlabs) reverse-biased by 64 DC voltage. The output current of the photodiode was electronically filtered (KR 2724, KR electronics), terminated with 50□Ω, and demodulated with a lock-in amplifier (HF2LI, Zurich Instruments) to achieve near shot-noise-limited sensitivity. The stimulated Raman loss signal at each pixel was sent to the analog interface box (FV10-ANALOG, Olympus) ofthe microscope to generate the image. All images were acquired with 80□μs time constant at the lock-in amplifier and 100□μs pixel dwell time (~27□s per frame of 512□×□512 pixels). Measured after the objectives, 40□mW pump power and 120□mW Stokes beam were used to image the carbon-deuterium 2183 cm^−1^ and off-resonance 2004 cm^−1^ channels.

### Phenazine detection by HPLC and spectrophotometry

10 μl of biofilm subcultures were spotted onto 4 mL of 1% tryptone 1% agar in 30 mm circular petri plates and grown for 72 hours at 25°C. The biofilm and agar were placed into 5 mL of 100% methanol and nutated overnight at room temperature to extract the phenazines. To measure the amount of aeruginosins, 600 μl of the methanol extract was added to 600 μl of chloroform, briefly vortexed, and allowed to settle into fractions. After separation, 100 μl of the top aqueous fraction was put into a 96-well plate and read in a plate reader. To quantify aeruginosin levels, the samples were excited at 520 nm and the emission read at 620 nm. The results presented are all normalized relative to the signal of PA14 biofilms. To measure levels of PCA, PCN, and PYO, the methanol extract was filtered through 0.22 μm cellulose acetate Spin-X centrifuge tube filters (Costar), then 200 μl loaded into HPLC vials for analysis. The resulting peaks were identified and quantified relative to samples of purified PCA, PCN, and PYO run in known concentrations.

### PMS sensitivity assay

Precultures of strains of interest were grown for 16 hours at 37°C, 200 rpm and then subcultured at a dilution of 30 μL into 3 mL LB medium in VWR 13 x 100 mm glass culture tubes. Subcultures were grown for about 2.5 hours to an OD (500 nm) of approximately 0.55, then normalized to an OD (500 nm) of 0.5 before serial dilution. 5 μl of each dilution was spotted onto 10 cm square plates containing 60 mL of 1% tryptone 1% agar medium with or without 800 μM PMS. Biofilms were incubated for 48 hours at 25°C before scanning on an Epson Expression 11000XL photo scanner. Brightness of all images was adjusted in the same way to a value of 150 using Adobe Photoshop for ease of visualization.

## Acknowledgments

This work was supported by NIH/NIAID grant R01AI103369 to L.E.P.D. and NIH/NIBIB grant R01EB029523 to W.M.

