## Supplementary material for "Spatial heterogeneity in biofilm metabolism elicited by local control of phenazine methylation": SI Figures 1-5

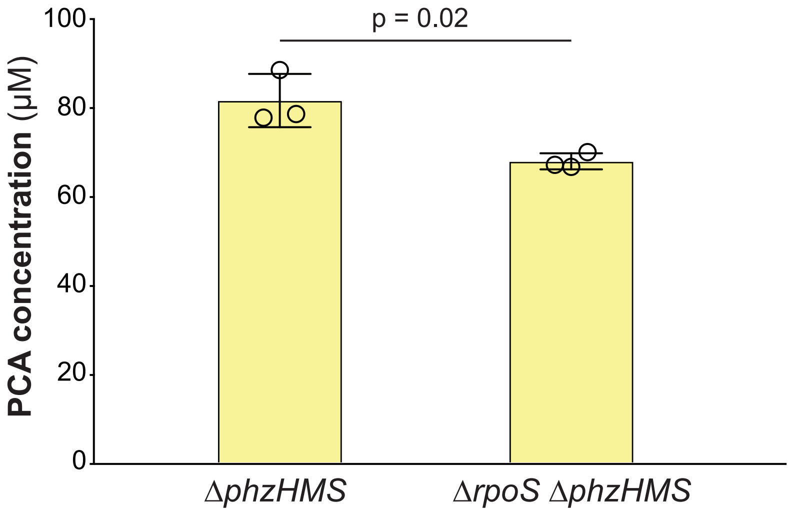


**Figure S1. *rpoS* deletion does not increase total phenazine production.** PCA produced by biofilms of PA14 and Δ*rpoS* strains lacking *phzH, phzM,* and *phzS* (Δ*phzHMS*). In these strains, the PCA production is indicative of the total amount of phenazines made due to the lack of the phenazine modification enzymes. Data points represent biological triplicates and error bars represent standard deviation. *p* values were calculated using unpaired, two-tailed *t*-tests.

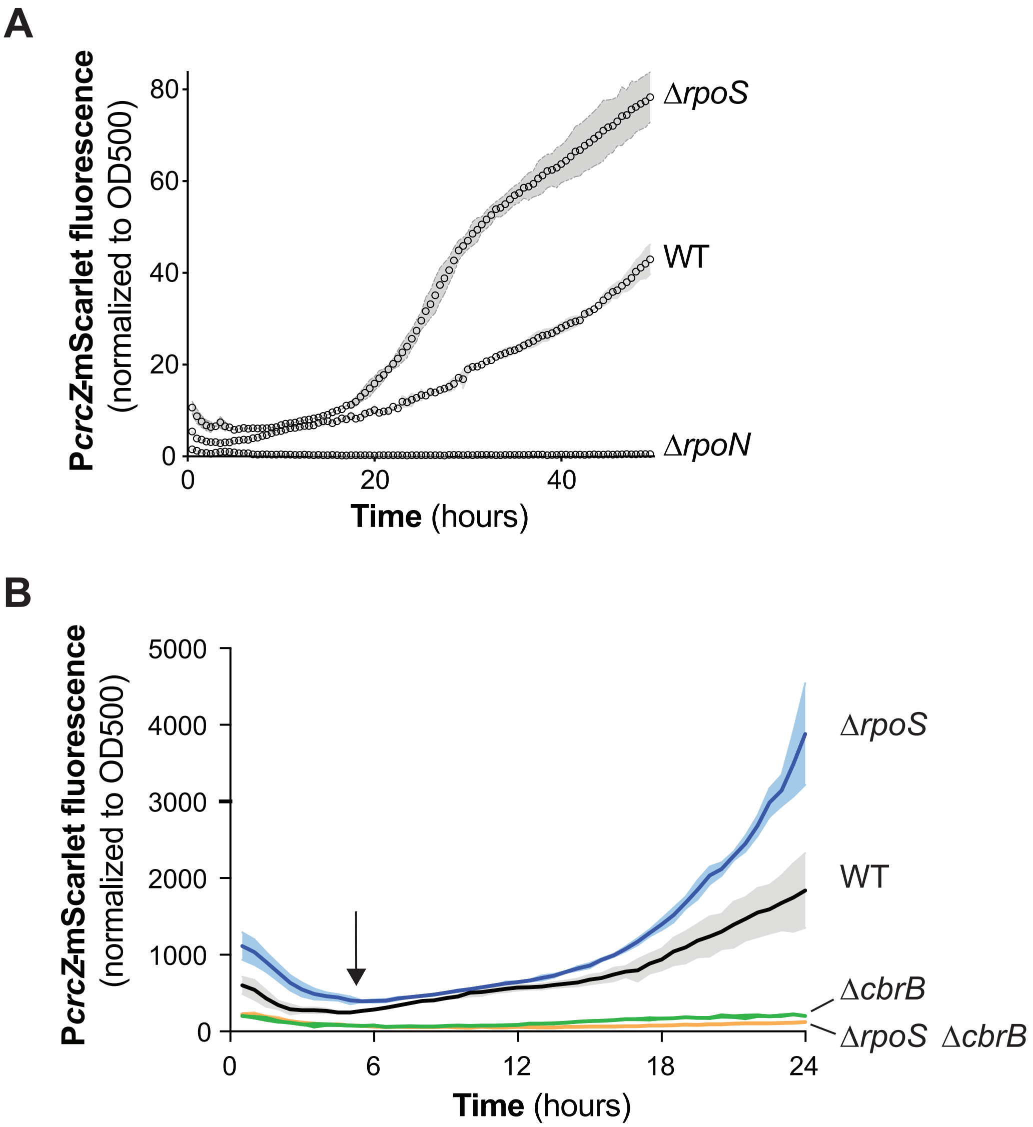


**Figure S2. Deletion of *cbrB* abrogates *crcZ* expression in the Δ*rpoS* background.** (**A**) PA14, Δ*rpoS*, Δ*rpoN* strains with P*crcZ-mScarlet* were grown in 1% tryptone broth with shaking for 24 hours. (**B**) PA14, Δ*rpoS,* Δ*cbrB,* and Δ*rpoS*Δ*cbrB* strains were grown in 1% tryptone broth with shaking for 24 hours. The arrow indicates the onset of stationary phase. Traces represent the averages of biological triplicates and shading indicates standard deviation.

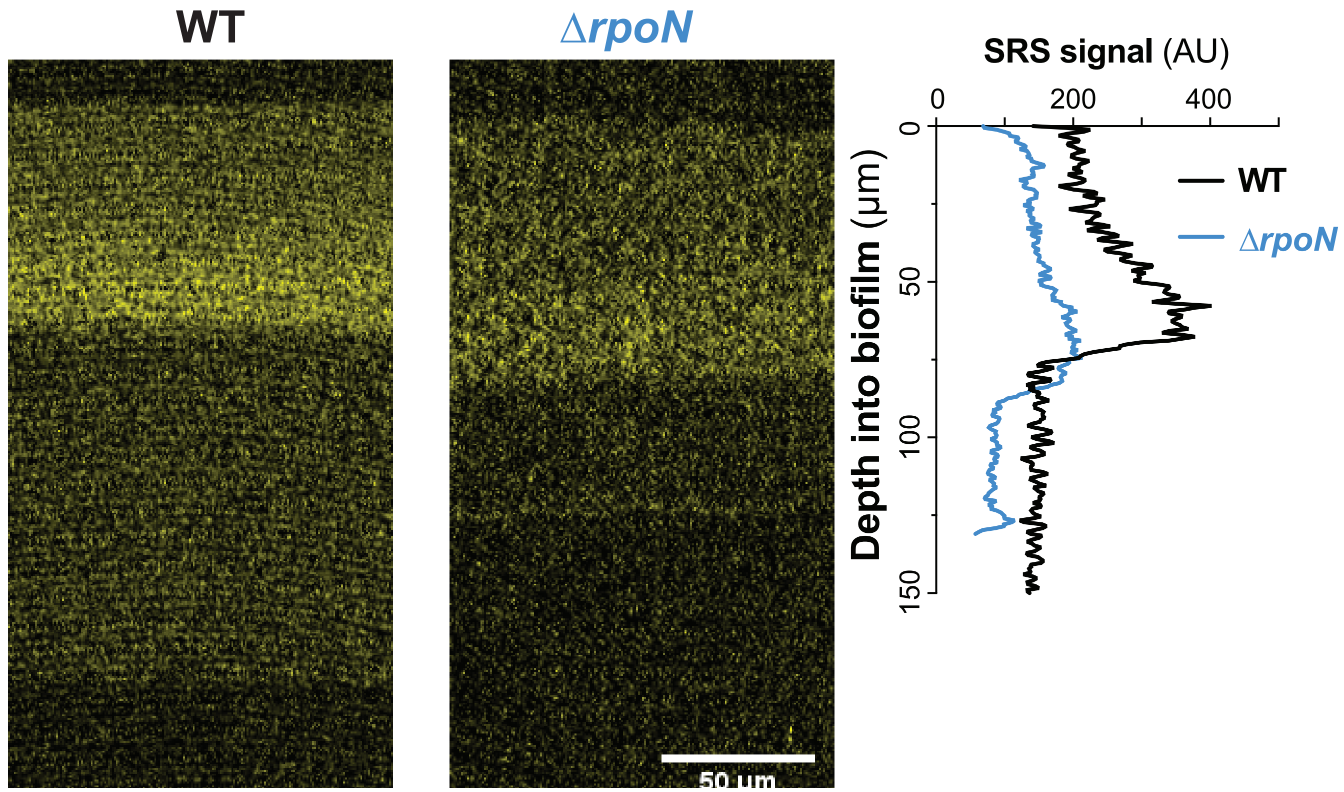


**Figure S3. The effect of RpoN on biofilm metabolic activity is consistent with its role in *crcZ* expression.** Left and center: SRS images of thin sections prepared from WT and Δ*rpoN* biofilms. SRS signal is indicative of metabolic activity and is false-colored yellow. Right: Average SRS signal across depth. The experiment was performed in biological triplicate and representative images are shown.

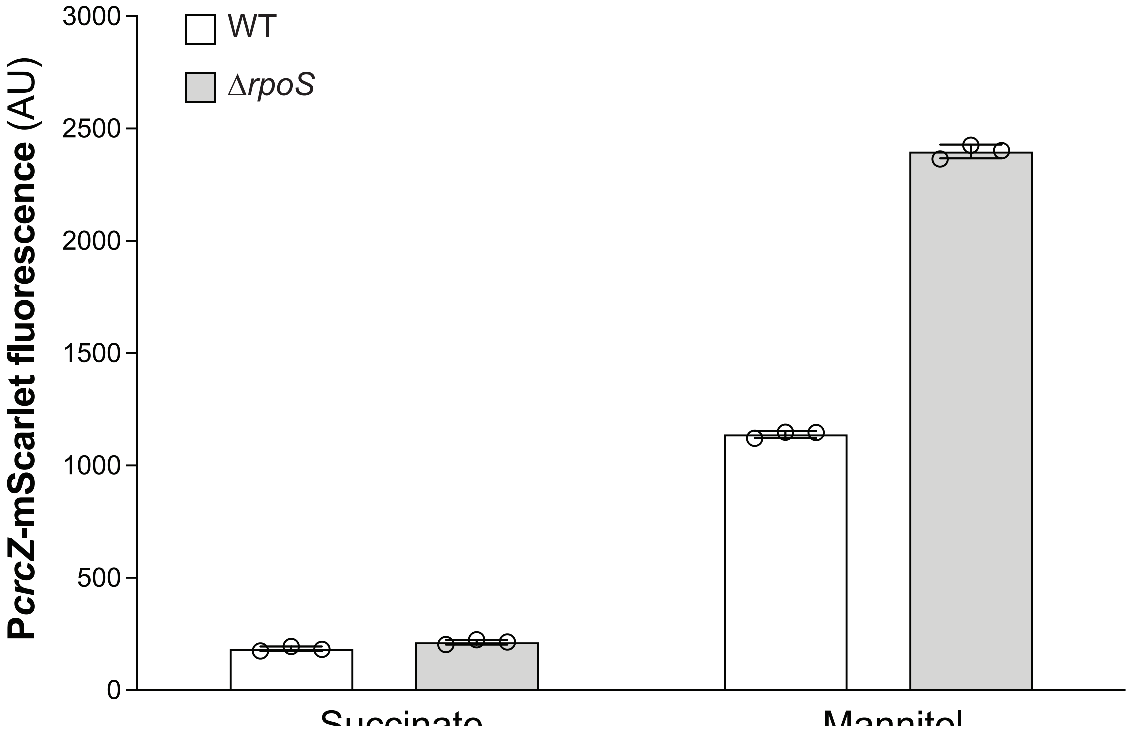


**Figure S4. *rpoS* deletion enhances *crcZ* expression during growth in liquid culture.** The WT and Δ*rpoS* P*crcZ-mScarlet* reporter strains were grown in a defined medium to an OD (500 nm) of 0.3 with the indicated compounds as sole carbon sources. Individual points represent biological triplicates and error bars represent standard deviation.

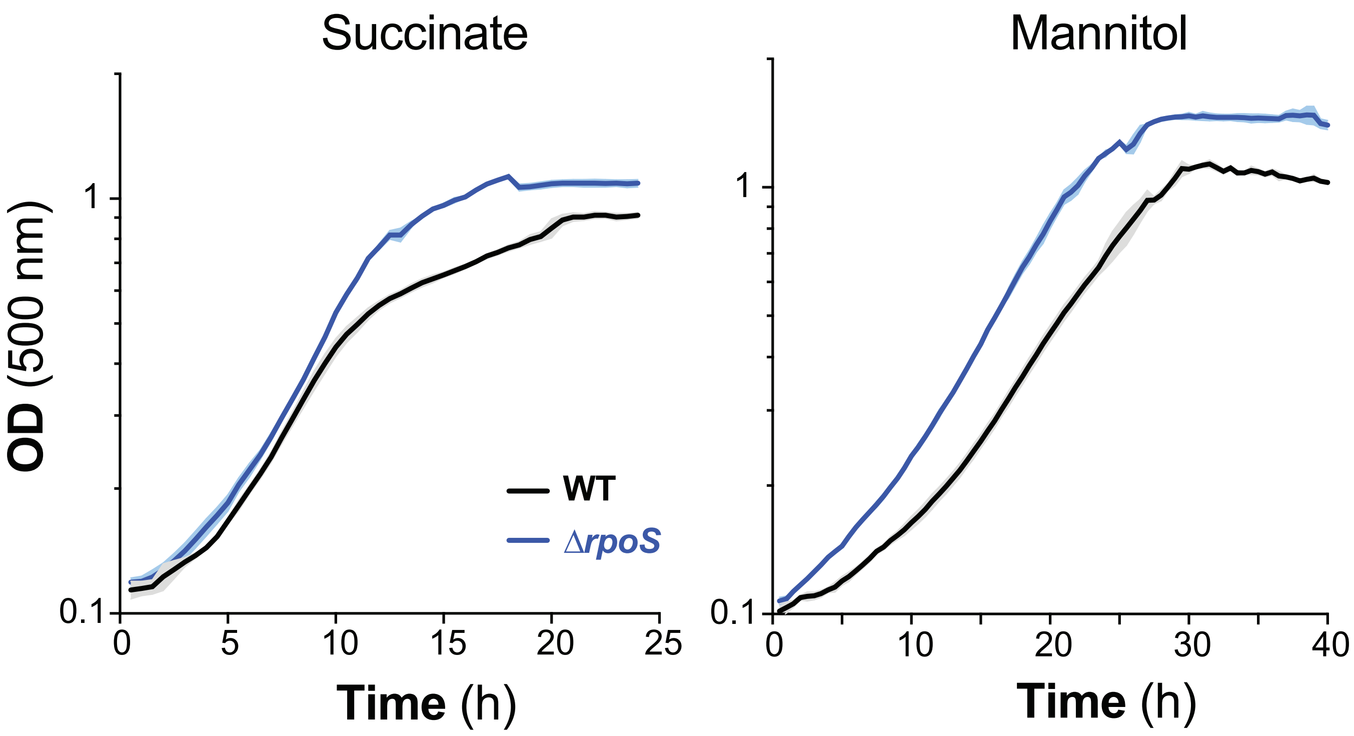


**Figure S5. Effects of *rpoS* and *crc* deletions on planktonic growth with individual carbon sources.** PA14 and Δ*rpoS* were grown in a defined medium with 20 mM succinate or mannitol. Traces represent the averages of biological triplicates and shading indicates standard deviation. Mannitol cultures were grown for 40 hours to capture the complete growth cycle.
