## Supplementary material for "Spatial heterogeneity in biofilm metabolism elicited by local control of phenazine methylation": SI Tables 1-3

**Table S1. Bacterial strains used in this study.**

| Number | Strain | Description | Source |
|---|---|---|---|
| *Pseudomonas aeruginosa* strains | | | |
| LD0 | UCBPP-PA14 (WT) | Clinical isolate UCBPP-PA14 | (1) |
| LD24 | Δ*phz* (also referred to as Δ*phz1/2*) | PA14 with the *phzA1-G1* (*PA14_09480-PA14_09410*) and *phzA2-G2* (*PA14_39970-PA14_39880*) operons deleted | (2) |
| LD3692 | Δ*phzH* | PA14 with *phzH* (*PA14_00640*) deleted | (3) |
| LD3739 | Δ*phzMS* | PA14 with *phzM* (*PA14_09490*) and *phzS* (*PA14_09400*) deleted | (3) |
| LD3746 | Δ*phzHMS* | PA14 with *phzH* (*PA14_00640*), *phzM* (*PA14_09490*), and *phzS* (*PA14_09400*) deleted | (4) |
| LD64 | BigBlue (*phzM+)* | DKN370; PA14 merodiploid strain containing an extra copy of *phzM* (*PA14_09490*) | (5) |
| LD3192 | Δ*rpoS* | PA14 with *rpoS* (*PA14_17480*) deleted | This study |
| LD3193 | Δ*rpoS*Δ*phz* | PA14 with *rpoS* (*PA14_17480*) and the *phz1* (*PA14_09480-PA14_09410*) and *phz2* (*PA14_39970-PA14_39880*) operons deleted | This study |
| LD3469 | Δ*rpoS*Δ*phzHMS* | PA14 with *rpoS* (*PA14_17480*), *phzH* (*PA14_00640*), *phzM* (*PA14_09490*), and *phzS* (*PA14_09400*) deleted | This study |
| LD3674 | Δ*crc* | PA14 with *crc* (*PA14_70390*) deleted | This study |
| LD3675 | Δ*crc*Δ*phz* | PA14 with *crc* (*PA14_70390*) and the *phz1* (*PA14_09480-PA14_09410*) and *phz2* (*PA14_39970-PA14_39880*) operons deleted | This study |
| LD3717 | Δ*rpoS*Δ*crc* | PA14 with *rpoS* (*PA14_17480*) and *crc* (*PA14_70390*) deleted | This study |
| LD3190 | Δ*rpoN* | PA14 with *rpoN* (*PA14_57940*) deleted | This study |
| LD3870 | PA14 attB::P*crc-mScarlet* | PA14 containing a construct in the *attB* site that expresses *mScarlet* under control of the 500bp region upstream of | This study |

| | | | |
|---|---|---|---|
| | | *crc* (*PA14_17480)* | |
| LD4082 | PA14 attB::P*crcZ-mScarlet* | PA14 containing a construct in the *attB* site that expresses *mScarlet* under control of the 350bp region upstream of *crcZ* (Unannotated between *PA14_62540* and *PA14_62560*. Annotated as *PA4726.11* in PAO1) | This study |
| LD3941 | Δ*rpoS* attB::P*crc-mScarlet* | Δ*rpoS* (*PA14_17480*) containing a construct in the *attB* site that expresses *mScarlet* under control of the 500bp region upstream of *crc* (*PA14_17480)* | This study |
| LD4108 | Δ*rpoS* attB::P*crcZ-mScarlet* | Δ*rpoS* (*PA14_17480*) containing a construct in the *attB* site that expresses *mScarlet* under control of the 350bp region upstream of *crc*Z (Annotated as *PA4726.11* in PAO1) | This study |
| LD4498 | Δ*cbrB* attB::P*crcZ-mScarlet* | PA14 attB::P*crcZ-mScarlet*, with *cbrB* (*PA14_62540)* deleted | This study |
| LD4501 | Δ*rpoS*Δ*cbrB* attB::P*crcZ-mScarlet* | PA14 attb::P*crcZ-mScarlet*, with *cbrB* (*PA14_62540)* and *rpoS* (*PA14_17480*) deleted | This study |
| LD4499 | PA14 attB::PTL*phzM-mScarlet* | PA14 containing a construct in the *attB* site in which the 695 bp upstream region and first 6 codons of *phzM* (*PA14_09490*) are fused to *mScarlet* | This study |
| LD4502 | Δ*rpoS* attB::PTL*phzM-mScarlet* | Δ*rpoS* (*PA14_17480*) containing a construct in the *attB* site in which the 695 bp upstream region and first 6 codons of *phzM* (*PA14_09490*) are fused to *mScarlet* | This study |
| LD4582 | Δ*crc* attB::PTL*phzM-mScarlet* | Δ*crc* (*PA14_17480)* a construct in the *attB* site in which the 695 bp upstream region and first 6 codons of *phzM* (*PA14_09490*) are fused to *mScarlet* | This study |
| *E. coli* strains | | | |
| LD44 | UQ950 | E. coli DH5α λ(pir) host for cloning; F-Δ(*argF-lac*)169 Φ80 d*lacZ58*(ΔM15) *glnV44*(AS) *rfbD1 gyrA96*(NalR) *recA1 endA1 spoT1 thi-1 hsdR17 deoR* λpir+ | D. Lies |
| LD661 | BW29427 | Donor strain for conjugation: *thrB1004 pro* | W. Metcalf |

| | | | |
|---|---|---|---|
| | | *thi rpsL hsdS lacZ* ΔM15RP4–1360 Δ(*araBAD*)567 Δ*dapA1341*::[*erm pir*(wt)] | |
| LD2901 | S17-1 | Donor strain for conjugation: $Str^R$, $Tp^R$, $F^-$ RP4-2-Tc::Mu *aphA*::Tn7 *recA* λpir lysogen | R. Simon |
| *Saccharomyces cerevisiae* strains | | | |
| LD676 | InvSc1 | *MATα/MATα leu2/leu2 trp1-289/trp1-289 ura3-52/ura3-52 his3-Δ1/his3-Δ1* | Invitrogen |

**Table S2. Plasmids used in this study.**

| Plasmid Name | Description | Source |
|---|---|---|
| pMQ30 | Yeast-based allelic-exchange vector; *sacB*$^+$, CEN/ARSH, URA3$^+$, $Gm^R$. | (6) |
| pFLP2 | Site-specific excision vector with cI857-controlled FLP recombinase. encoding sequence, sacB$^+$, $Amp^R$. Used to insert LD3208-based plasmids into P. aeruginosa strains. | (7) |
| pLD3208 | $Gm^R$, $Tet^R$ flanked by Flp recombinase target (FRT) sites to resolve out resistance cassettes. | (8) |
| pLD3471 | ∆*rpoS* (*PA14_17480*) PCR fragment introduced into pMQ30 by gap repair cloning in yeast strain InvSc1. | This study |
| pLD3473 | ∆*rpoN* (*PA14_57940*) PCR fragment introduced into pMQ30 by gap repair cloning in yeast strain InvSc1. | This study |
| pLD3673 | ∆*crc* (*PA14_70390*) PCR fragment introduced into pMQ30 by gap repair cloning in yeast strain InvSc1. | This study |
| pLD3869 | 500 bp upstream of *crc* (*PA14_70390*) PCR fragment ligated into pLD3208 using SpeI and XhoI. | This study |
| pLD4645 | 350 bp upstream of *crcZ* (annotated as *PA4726.11* in PAO1) PCR fragment ligated into pLD3208 using SpeI and XhoI. | This study |
| pLD4179 | 695 bp upstream and first six codons of *phzM* (*PA14_09490*) PCR fragment ligated into pLD3208 using EcoRI and SphI. | This study |

**Table S3. Primers used in this study.**

| Primer Number | Sequence |
|---|---|
| Primers for plasmid pLD3471 (used to make ∆*rpoS*) | |
| LD2560 | ggaattgtgagcggataacaatttcacacaggaaacagct TGGATAAGGGGGAAGGATTG |
| LD2561 | CCGTTCTTCTCCAGGATCTC CGGCCCTTCTTTTTTGAGTGC |
| LD2562 | GCACTCAAAAAAGAAGGGCCG GAGATCCTGGAGAAGAACGG |
| LD2563 | aggcaaattctgttttatcagaccgcttctgcgttctgat AAACCACCAGCCTGCCGCAC |
| Primers for plasmid pLD3473 (used to make ∆*rpoN*) | |
| LD2568 | ggaattgtgagcggataacaatttcacacaggaaacagct CGCGCCCGCGCATCGACATG |
| LD2569 | CACCAGTCGCTTGCGCTC CATCTTGAGGACTAGCGATGG |
| LD2570 | CCATCGCTAGTCCTCAAGATG GAGCGCAAGCGACTGGTG |
| LD2571 | aggcaaattctgttttatcagaccgcttctgcgttctgat CAGGGCGCGCTGCGCCAGGT |
| Primers for plasmid pLD3673 (used to make ∆*crc*) | |
| LD3184 | ggaattgtgagcggataacaatttcacacaggaaacagct GCCCTTGTCGTTGACGTAGC |
| LD3185 | TCGACGATCAGCGGCGCATGC CCGCAGCCTGAATACCATTCAC |
| LD3186 | GTGAATGGTATTCAGGCTGCG GCATGCGCCGCTGATCGTCGA |
| LD3197 | aggcaaattctgttttatcagaccgcttctgcgttctgatTCGGCGAGAACACCCTGTAC |
| Primers for plasmid pLD3869 (used to make *Pcrc-mScarlet*) | |
| LD3273 | tcccgacgggcccggtaccaGATGATCTGCATCACTTCG |
| LD3274 | tcttaaatctagactcgaggAAATGGCCCCCAAAATCAC |
| Primers for plasmid pLD4645 (used to make *PcrcZ-mScarlet*) | |
| LD3663 | acgtacactagtCACCCTGCAACCTGTTACC |
| LD3272 | tcttaaatctagactcgaggCAATACATAAGCAGATGCCGTGCC |
| Primers for plasmid pLD4179 (used to make *PTLphzM-mScarlet*) | |
| LD3975 | acgtacgtacGAATTCGCGCCGCCTCCGAGA |
| LD3660 | ctcctttactaagattcgaattattcatcttttattc |

| | |
|---|---|
| LD3751 | ttcgaatcttagtaaaggagaagctgtg |
| LD3976 | acgtacgtacGCATGCccagtgcaggagctcataaaac |
